# Substitution of Lead Tungstate for Lead Abellaite in a Nanoparticle-Alginate Nanocomposite as a Contrast Agent for Post-Mortem CT Imaging: *In Vitro* Bulk Performance Evaluation

**DOI:** 10.64898/2026.02.27.708615

**Authors:** Anna X Le, Steven W. Buckner, Paul A. Jelliss, Sara McBride-Gagyi

**Affiliations:** The Ohio State University, Department of Biomedical Engineering, Columbus, OH; Saint Louis University, Department of Chemistry, Saint Louis, MO; The Ohio State University, Department of Orthopaedic Surgery, Columbus, OH

## Abstract

Accurately assessing blood vascular networks is important in all organ systems in health, disease, and healing. However, methods to do so in a holistic fashion in post-mortem specimens have significant limitations. We have previously demonstrated proof of concept for a lead abbelaite nanoparticle and alginate nanocomposite contrast agent which would allow greatly improved vessel imaging using x-ray based imaging modalities like CT and microCT. Specifically the contrast is spectrally enhanced to easily allow segmentation from mineralized tissues and gelation is triggerable permitting better vascular perfusion. Here we expand upon that work by substituting lead tungstate nanoparticles. We found that delivery viscosity and radiopacity are largely unaffected. However, mechanical strength was negatively impacted as abellaite presence was lowered. In sum, these formulations have performed in bulk reasonably enough to warrant advancement to in vivo post-mortem evaluation in small animal models.

## Introduction

Assessing the location, connectivity, and structural characteristics of cardiovascular vessel networks is critical for understanding disease progression, tissue remodeling, and injury repair.^1^ Histology remains a gold-standard technique for resolving small biological features such as microvasculature and individual cells, but it is inherently invasive, labor-intensive, and destructive. In addition, histological sectioning introduces sampling and alignment biases that can significantly affect quantitative and spatial analyses. Computed tomography (CT) and microCT are non-destructive, x-ray–based imaging techniques that enable three-dimensional visualization of intact biological specimens with relatively few contraindications.^2–5^ Owing to their high spatial resolution, these modalities more accurately capture the tissues’ spatial organization and distribution across multiple length scales compared to typical magnetic resonance imaging (MRI) or positron emission tomography (PET).^6–8^ CT and microCT are also among the most cost-effective advanced imaging techniques, supporting their widespread use in clinical diagnostics, preclinical research, and post-mortem studies.^7,8^ However, because most soft tissues exhibit similar x-ray attenuation coefficients, CT and microCT alone provide limited intrinsic soft-tissue contrast, necessitating the use of radiopaque contrast agents to differentiate structures such as vasculature, muscle, and connective tissue. ^5,9–13^

Several CT and microCT vascular contrast agents have been developed, yet the optimal material properties depend strongly on the intended application.^5,12,13^ Across all use cases, high radiopacity and efficient vascular perfusion, including capillaries with diameters below 5 µm, are desirable. *In vivo* and live *ex vivo* contrast agents must clear from the system within hours to days while minimizing toxicity and physiological disruption. In contrast, post-mortem contrast agents must remain stable for days to months to withstand tissue harvesting, handling, storage, and downstream processing without significant loss of radiopacity or structural fidelity until scanning. Currently available cost-effective products for post-mortem applications have several limitations such as poor vascular perfusion, upstream pressure-induced damage or rupture, and similar radiopacity to bone ^11^.

In our previous work, we reported an alginate (ALG)-based post-mortem vascular contrast agent incorporating abellaite (AB, lead carbonate, NaPb_2_(CO_3_)_2_OH, ρ = 5.93 g/cm^3^, crystallite size 22 ± 1 nm) nanoparticles (NPs) as both the contrast and alginate-crosslinking element ^14,15^. In theory this novel system should not spontaneously gel until the pH is moderated lowered (<6.5), which was supported by limited proof-of-concept testing ^14,15^. We used d-(+)-gluconic acid δ-lactone (GDL) as the pH-modifying trigger. At AB concentrations of 0.1 g/mL and 0.4 g/mL in 1% ALG the hydrogels were similar with respect to initial viscosity, gelation times, and mechanical strength. Nanocomposites gelled in approximately 10 to 18 min after GDL addition depending on concentration while mechanical behavior at physiological and processing strains (< 25%) was unaffected. On the other hand, radiopacity at 0.1 g/mL was not reliably more radiopaque than bone. Our lab’s primary interest in vascular structure is the evolution of angiogenesis in repairing bone. Sufficiently high spectral contrast is needed to separate the contrast-filled vessels from the bone tissue in a single microCT scan. Vascular network research in other organ systems or processes may not need the same spectral differences but would still benefit high contrast agent radiopacity. Increasing the concentration to 0.2 g/mL and above consistently reached the maximal level of detection at typically scanning conditions^14^. While this highly informative study demonstrated the feasibility of combining radiopaque nanoparticles and ALG for post-mortem microCT angiography, additional studies and refinements are needed to investigate spontaneous gelation in the absence of the GDL trigger and better characterize the 0.2 g/mL formulation behaviors.

Further, it could be advantageous to incorporate a more compact and less soluble nanoparticle to enhance radiopacity while limiting off target effects. Lead tungstate is an ideal candidate (LT, PbWO_4_, ρ = 8.28 g/cm^3^). At estimated scanning energies of 30 keV, individual AB and LT molecules should have equivocal radiopacity ^16^. However, LT’s higher density should result in more molecules per unit weight of NP-powder resulting in a higher overall radiopacity. Also, tungstate’s more basic disassociation constant (pKb ∼7.5 vs carbonate’s pKb ∼3.7 (XX)) should keep these molecules from dissolving while the ABNPs disassociate and initiate ALG crosslinking. This would limit potential leaching into surrounding tissues and increase safety for users. However, having fewer ABNPs could compromise overall gel mechanical stability and strength.

Building on our previous work, the present study investigates various concentrations of AB and LT NPs embedded within an ALG matrix. We hypothesize that none of our formations will spontaneously gel within anticipated perfusion durations. Further, increasing LT subsitution will moderately increase overall radiopacity, as LT contains both lead and tungsten in a more compact molecule,^9^ but eventually compromise hydrogel material strength.

## Materials & Methods

### LTNP Synthesis & Characterization

[TBA]

### ALG-NP Formulations

ALG [Alginic acid sodium salt, 350 to 550 mPa.s (1% at 20°C,Brookfield LV), ThermoFisher Scientific, Catalog number 177772500] was used as the hydrogel matrix carrier, and GDL (Sigma-Aldrich, Catalog Number G4750) served as a gelation trigger agent. Calcium chloride (CaCl_2_, Sigma-Aldrich, Catalog Number 223506) was used in select experiments to simulate physiological calcium exposure. All reagents were used as received unless otherwise specified.

All formulations utilized 1% (w/v) ALG dissolved in deionized (DI) water, prepared by dissolving 0.50 g of alginic acid sodium salt per 50 mL of final total volume. NP–ALG suspensions were prepared by adding either 2.0 g or 4.0 g of NP powder per 10 mL of solution to achieve final nanoparticle concentrations of 0.20 or 0.40 g/mL, respectively. NP powders containing AB, LT, or combinations of both nanoparticle types were evaluated. The formulations described in Table 1 were used for all experiments. A 0.4g/mL 0:1 formulation was originally included, but it spontaneously gelled immediately upon combination. Thus LT-only gels were not deemed worth further investigation.

**Table 1:** NP-ALG Formulations Tested.

| Concentration (g/mL) | Ratio | LT Mass (g/mL) | AB Mass (g/mL) |
| --- | --- | --- | --- |
| 0.4 | 0:1 | 0 | 0.4 |
|  | 1:1 | 0.2 | 0.2 |
|  | 2:1 | 0.27 | 0.13 |
|  | 5:1 | 0.33 | 0.07 |
| 0.2 | 0:1 | 0 | 0.2 |
|  | 1:1 | 0.1 | 0.1 |
|  | 2:1 | 0.13 | 0.07 |

### Initial Viscosity and Rheological Characterization

Rheological measurements (n=3/formulation) were performed on NP–ALG formulations to determine the initial viscosity and detect any spontaneous gelation of the nanocomposite hydrogel precursor solutions without GDL addition. NP powders were added to the ALG solution and sonicated on ice for approximately 30 seconds to ensure homogeneous distribution and immediately subjected to oscillatory shear testing for 1800 sec (30 min) using CP1/40 SR0089 TI cone geometry (25°C, constant strain amplitude of 1%, 1.6 Hz, Kinexus rotational rheometer). The loss modulus (G’’), storge modulus (G’), tan(δ), and complex viscosity (η*) were recorded throughout the test period.

### Compressive Mechanical Testing

Unconfined compression testing was conducted to evaluate whether the solidified gels could tolerate postmortem tissue extraction and routine handling prior to microCT scanning. Cylindrical specimens (12.7 mm diameter × 12.7 mm height; 2–5 samples per batch, 1–2 batches per formulation; total n = 3–5 per group) were fabricated from NP–ALG–GDL hydrogels. GDL powder (0.06 g/mL GDL) was added to the sonicated NP-ALG solution and mixed just prior to casting. Custom molds were fabricated by plasma bonding silicone wells onto glass microscope slides, producing specimens with silicone sidewalls and a glass base. After filling the wells, a second glass slide was placed atop each mold to ensure parallel, flat upper and lower surfaces. Within each batch, half of the specimens were carefully demolded and tested immediately. The remaining samples were demolded, immersed in a 2 mM CaCl_2_ solution for 1 hour to mimicking potential secondary ionic cross-linking from surrounding tissues *in vivo*, and subsequently tested.

Specimens were compressed to failure (10.0 mm/min, Instron Electropuls E3000 with 450 N load cell). At each test start, the specimens were not in contact with the top platen so that specimen height could be accurately determined from the loading curve. A custom MATLAB script was used to analyze the resulting force-displacement curves and transform them into stress-strain curves. These curves were used to determine gel specimen height and the following mechanical parameters: including the initial elastic modulus (from the linear region of the stress– strain response), stress at 10% and 25% strain, failure strain, failure stress, and material toughness.

### Long-term Stability

To evaluate long-term structural stability, additional matched specimens were prepared using the same solution preparation, casting, and, if applicable, CaCl_2_ immersion methods described for compression testing. These were stored in humidity chambers at room temperature for several weeks. Structural integrity and macroscopic changes were documented through periodic imaging.

### Radiopacity Assessment

The intended imaging application, microCT, was used to evaluate radiopacity in comparison to bone. NP–ALG–GDL solutions were made as described for compression testing. Mixed solutions were cast within small-diameter tubing (0.38 mm inner diameter; Tygon S3) and allowed 30 min to preliminarily solidify. The filled tubing was sectioned into approximately 10 mm-long segments (3 samples per batch, 3 independent batches; total n = 9 per group). For these experiments all specimens were immersed in CaCl_2_ for 1 hour. Each hydrogel-filled segment was positioned within a custom-designed, 3D-printed sample holder alongside a mouse femur and a potassium phosphate (K_2_HPO_4_, 1,000 mg/mL) calibration phantom for reference. Throughout the remaining storage and micro-computed tomography (microCT) imaging, the tubing samples and holders were fully submerged in a 2 mM CaCl_2_ solution to preserve hydration and promote long-term structural stability. Samples were then scanned (70 kV, 114 μA, 8 W, integration time 1000 ms, 15 μm resolution, ScanCo uCT50). Reconstructed images were analyzed to assess x-ray attenuation and contrast uniformity (Dragonfly ORS). Image intensity values were calibrated two ways to address the different units used by clinicians, Houndsfeld Units (HU), and small animal imaging systems, mgHA/cm^3^. The surrounding storage medium was set at zero in both unit systems. For HU scaling, the other reference point, air, was set at -1000 HU. For mgHA/cm^3^ scaling, the potassium phosphate phantom served as the other reference point at 1,000 mgHA/cm^3^. Segmentation masks were generated for each hydrogel sample, the femur bone tissue material, and the calibration phantom. Scaled voxel intensities values were calculated for each mask.

### Statistics

For rheological testing, a repeated-measures one-way analysis of variance (ANOVA) was conducted to evaluate time-dependent changes in complex viscosity within each formulation. This approach was used to identify the time point at which viscosity values became significantly different from baseline. Compression data were analyzed using two approaches. First, data from the seven NP–ALG groups were evaluated using multifactor ANOVA to determine the independent and interactive effects of nanoparticle concentration, AB:LT mass ratio, and CaCl_2_ exposure on mechanical outcomes. Second, mechanical data from the seven NP–ALG formulations and two historical control materials (12% gelatin and Microfil) were analyzed using one-way ANOVA with group as the independent factor. This historical control data was collected using identical methods for a prior study. This analysis was performed to assess overall differences among all groups, including comparisons with historical controls, and to support subsequent pairwise comparisons within the NP–ALG formulations. MicroCT-derived radiopacity measurements from the seven NP–ALG formulations and two control groups were similarly analyzed using one-way ANOVA (factor:group). All statistical tests were performed using MATLAB. For all ANOVA models, statistically significant main effects (p < 0.05) were followed by appropriate post-hoc pairwise comparisons between relevant groups. Statistical significance was defined as p < 0.05.

## Results

### Rheological Characterization

Rheological characterization was performed to assess the initial viscosity and flow stability of alginate–nanoparticle formulations in the absence of GDL-induced gelation. Across all formulations, complex viscosity increased gradually over the 30-minute testing period but remained low, indicating stable, fluid-like behavior without evidence of spontaneous or shear-induced gelation (Figure 1). Alginate alone exhibited an approximately 10% increase in viscosity over 30 minutes (p = 0.002), while formulations containing nanoparticles showed comparable viscosity increases ranging from 8–20% over the same period. Formulations containing 0.2 g/mL total nanoparticle concentration exhibited consistently lower viscosities than those at 0.4 g/mL, while 1% alginate alone displayed the lowest viscosity throughout testing. Notably, the 0.4 g/mL alginate–nanoparticle formulations demonstrated a significant increase in viscosity after approximately 14.5 minutes, whereas the 0.2 g/mL formulations showed no time-dependent viscosity increase beyond baseline trends. Importantly, even at the higher nanoparticle concentration, viscosities remained within a range compatible with vascular perfusion, suggesting minimal risk of premature flow restriction or vessel blockage prior to intentional gelation. Although high lead tungstate substitution increased viscosity by approximately 70% (p < 0.001), all formulations remained injectable.

**Figure 1:**
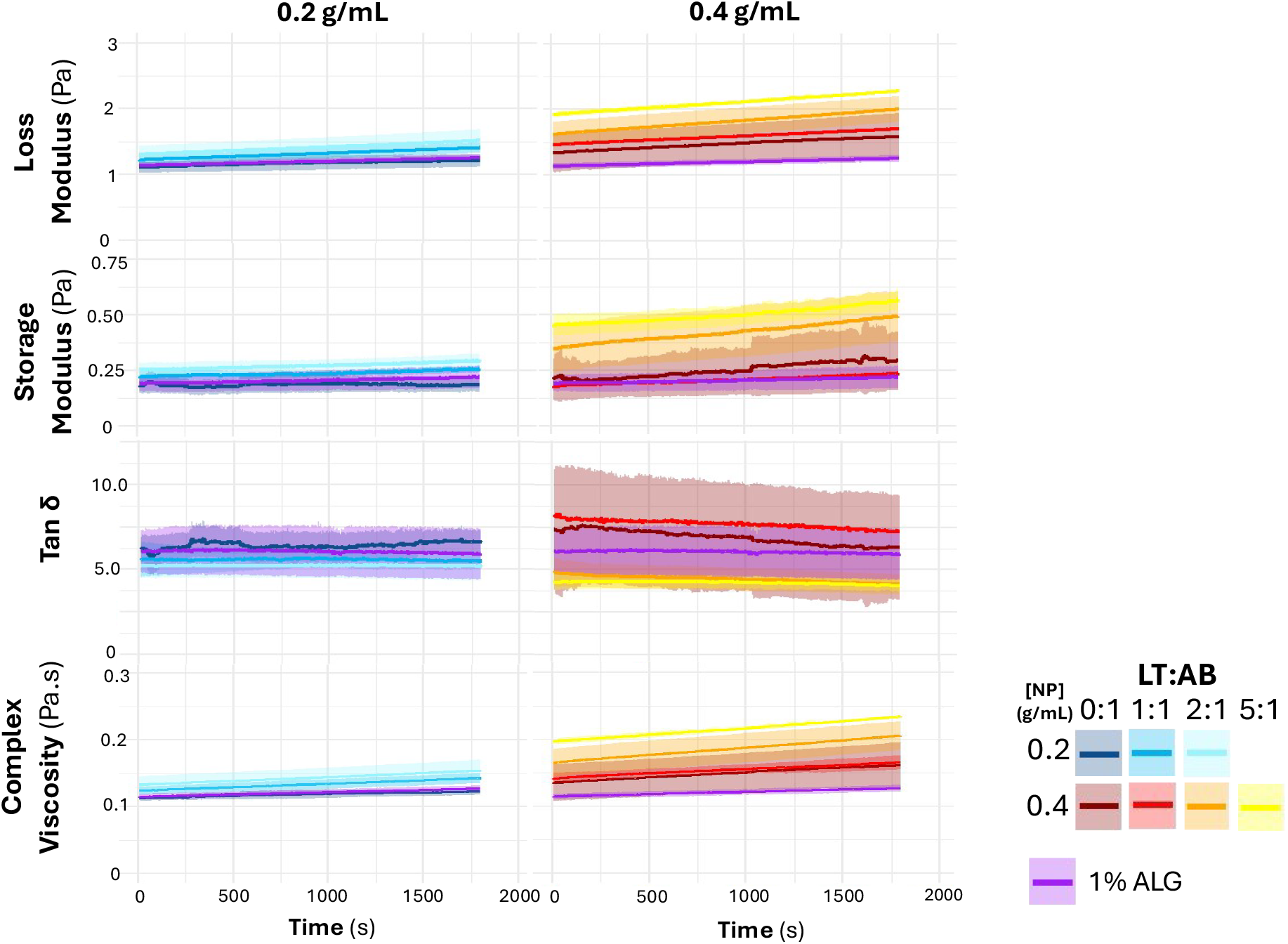
Results under Oscillatory Shear. Time-dependent changes in storage modulus, loss modulus, complex viscosity (η*), and tanδ for 0.20 g/mL (left panels) and 0.40 g/mL (right panels) NP–ALG formulations. For both concentrations, moduli and complex viscosity increased gradually over time, with the 0.40 g/mL formulations exhibiting greater magnitude changes compared to 0.20 g/mL. Tanδ showed minor temporal variation but remained relatively stable overall. Increasing the LT:AB mass ratio (greater LT substitution) resulted in higher moduli and viscosity, whereas tan δ demonstrated an inverse trend.

Statistical analysis revealed that time had a significant effect on viscosity (p < 0.001), with changes becoming apparent after approximately 245 s, consistent with slow structural evolution rather than abrupt network formation. Lead tungstate concentration was also a significant contributor to viscosity (p < 0.001), and a significant interaction between lead tungstate and abellaite concentrations was observed (p = 0.032). In contrast, abellaite concentration alone did not significantly influence viscosity (p = 0.07). Collectively, these results demonstrate that nanoparticle loading and composition modulate viscosity in a predictable manner, while all formulations remain sufficiently fluid over clinically relevant time scales. This rheological stability supports their suitability for vascular perfusion applications, ensuring adequate working time prior to controlled gelation without inducing shear-related clogging or premature solidification.

### Compressive Mechanical Testing

Unconfined compression testing was performed to evaluate the mechanical behavior of gelled nanoparticle–alginate formulations, with and without post-gelation calcium exposure, and compared against historical 12% gelatin and Microfil controls. Representative stress–strain curves for all formulations are shown in Figure 2. Across all conditions, gels exhibited nonlinear elastic behavior typical of ionically crosslinked hydrogels.

**Figure 2:**
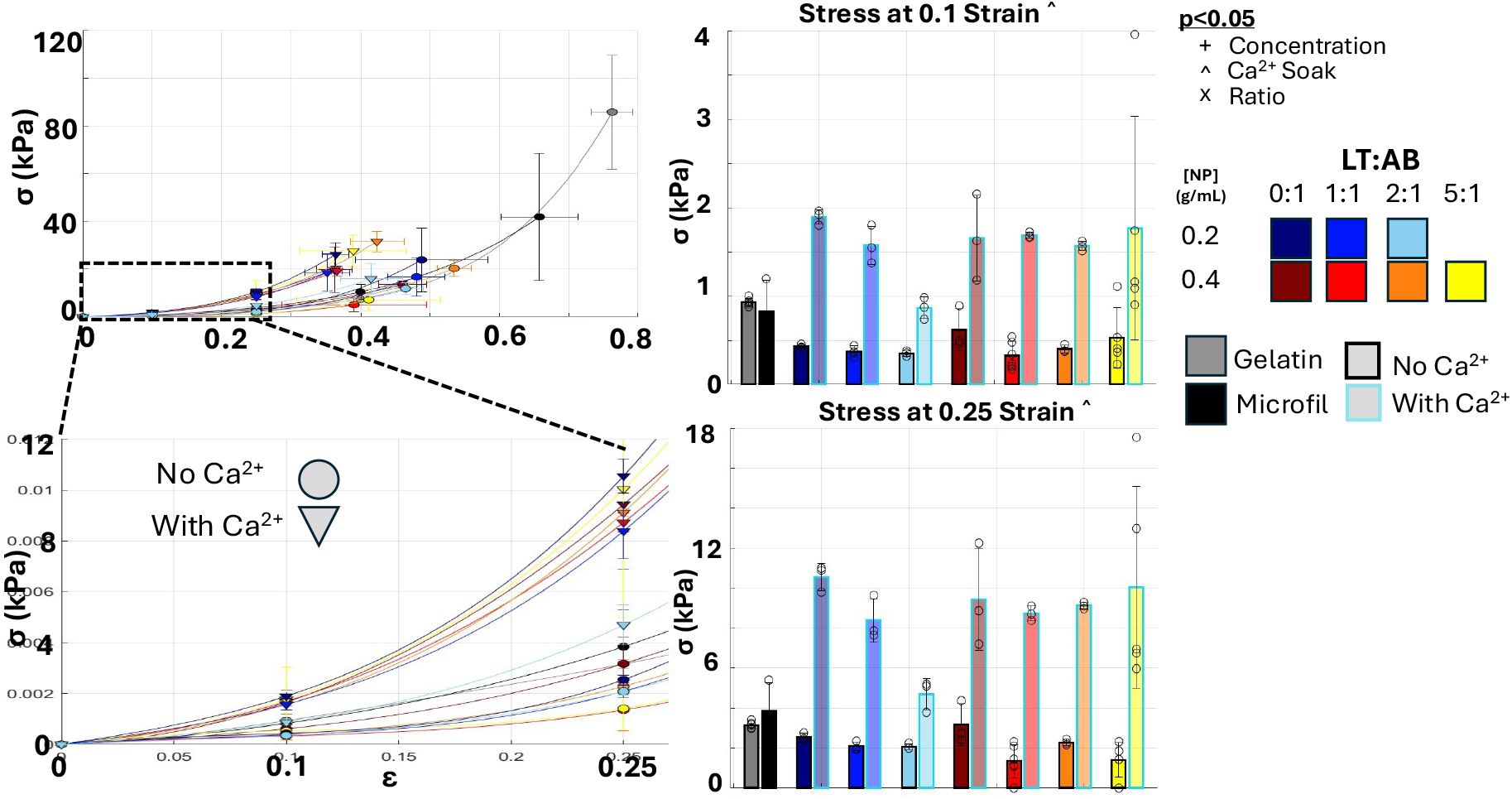
Stress-Strain Curves for Controls and NP-ALG gels. (Top panel) Full stress–strain curves for all experimental and control groups under unconfined compression. (Bottom panel) Enlarged view of the low-strain region corresponding to deformation levels relevant to tissue harvesting and handling.

Control materials, gelatin and Microfil, tolerated higher overall strains prior to failure; however, within the low-strain operating range relevant for vascular perfusion and structural support (ε ≤ 0.25), nanoparticle–alginate formulations demonstrated comparable or superior stress responses. In particular, calcium-soaked gels consistently generated higher stresses than their unsoaked counterparts, indicating increased stiffness following calcium exposure. This effect was evident across all nanoparticle concentrations and AB:LT ratios.

At 0.1 and 0.25 strain, stress values were quantified and summarized in bar plots (Figure 2B). At both strain levels, calcium-soaked samples exhibited significantly greater stress than unsoaked gels as well as both control materials. Notably, at 0.25 strain, all calcium-treated formulations showed a pronounced upward shift in stress compared to their corresponding GDL-only conditions, highlighting the reinforcing effect of calcium-mediated crosslinking. In contrast, unsoaked gels displayed lower stress values and greater variability, particularly at higher nanoparticle ratios.

Statistical analysis using two-way ANOVA identified post-gelation calcium soaking as the dominant determinant of compressive mechanical performance across all formulations (Figure 3). Calcium exposure produced a coordinated shift in mechanical behavior, significantly increasing elastic modulus, toughness, gauge length at failure, stress at low strains, and stress at failure (p < 0.0001 for all), while simultaneously reducing failure strain (p < 0.0016). Together, these changes indicate a transition toward stiffer, stronger, and more mechanically robust gels with reduced extensibility, consistent with enhanced ionic crosslinking following calcium treatment.

**Figure 3:**
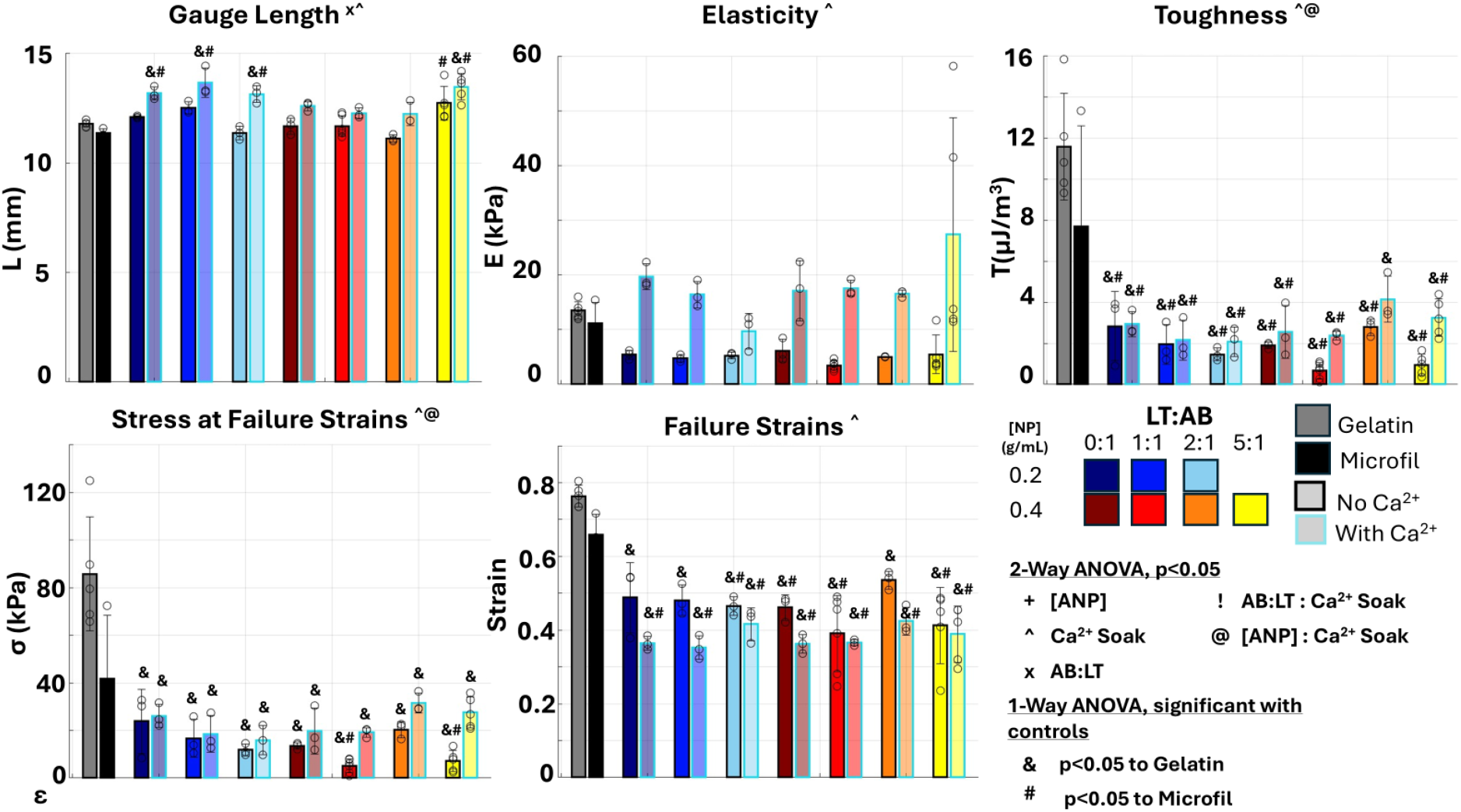
Mechanical Properties of Controls and NP-ALG Gels.

**Figure 4:**
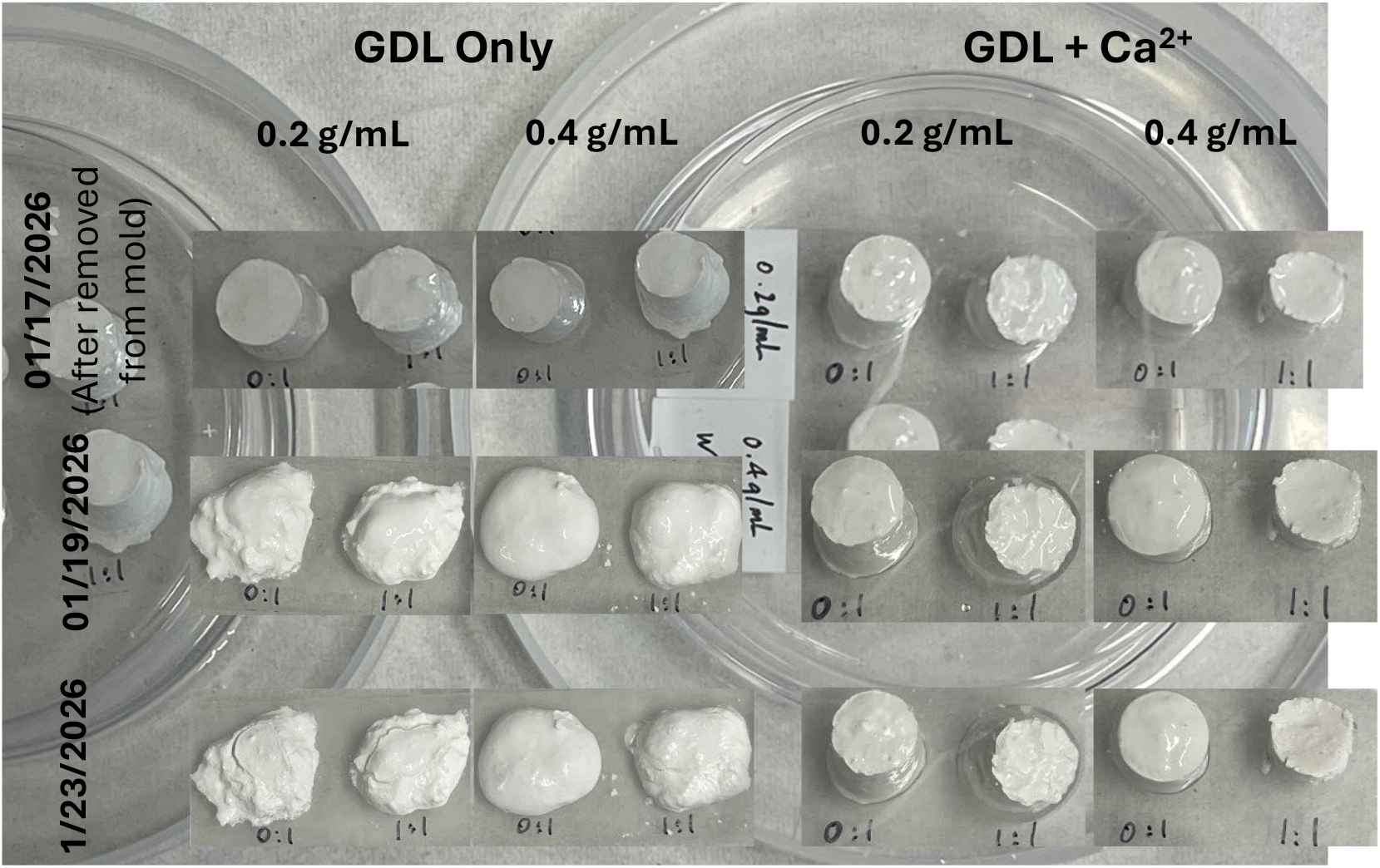
Calcium Soaking for Structural Stability. Images of 0.20 and 0.40 g/mL NP–ALG hydrogels with varying LT:AB ratios from matched batches. Samples exposed to short-term soaking in physiologically relevant calcium concentrations maintained structural integrity over extended storage, with no observable macroscopic degradation over time.

**Figure 5:**
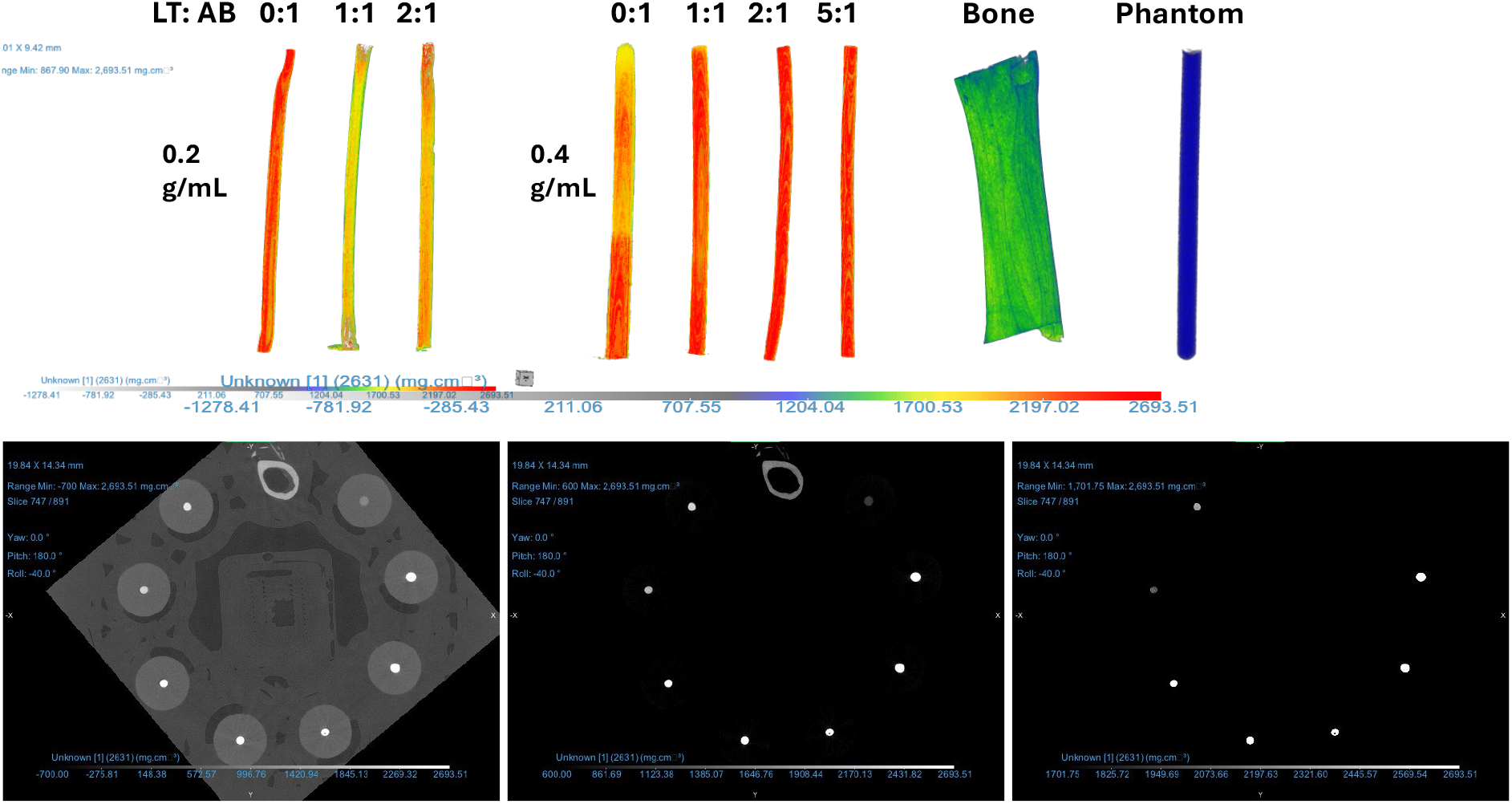
MicroCT images of Controls and Gels. (A) 3D images for controls and NP-ALG gels with the color scale bar shows the uniform gels with no bubbles or deformations while storing. (B) The same microCT slice visualized with low, medium, and high threshold are used. With high threshold, the bones and phantom can easily be segmented from the gels regardless of formulations.

**Figure 6:**
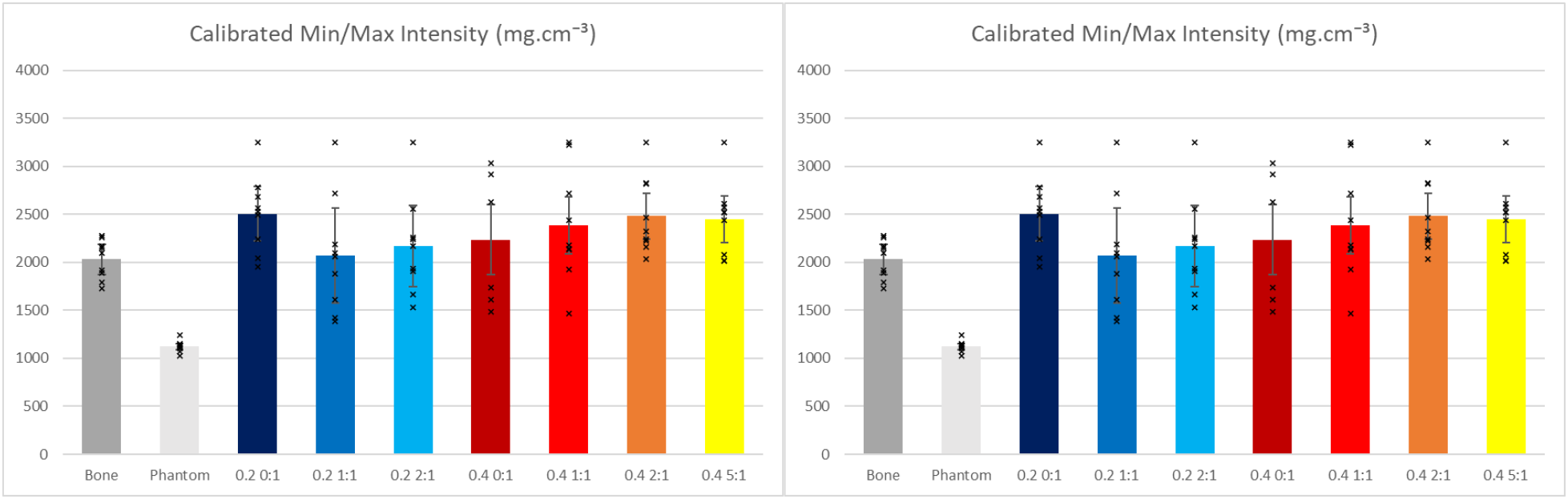
Mineral Density of Bone, Phantom, and NP-ALG gels. All

In contrast, nanoparticle composition played a secondary, modulatory role. The AB:LT ratio significantly influenced gauge length (p = 0.0161) and stress at failure but did not significantly affect elastic modulus or stress at low strains, indicating that nanoparticle identity primarily alters failure behavior rather than bulk stiffness. Nanoparticle concentration alone did not produce statistically significant changes in any measured mechanical metric, underscoring that mechanical reinforcement arises primarily from calcium-mediated network strengthening rather than increased particle loading.

### Structural Stability

Long-term structural stability was preliminarily evaluated to determine whether the gel formulations obviously degraded over time. All gels initially appeared macroscopically intact regardless of calcium treatment, exhibiting well-defined cylindrical geometry. However, pronounced differences emerged over time. Gels prepared without calcium exposure progressively lost structural integrity, with visible softening and eventual reliquification.

In contrast, calcium-soaked gels remained dimensionally stable throughout the entire 18-day observation period. These samples retained their original shape and mechanical coherence, with no evidence of liquefaction or gross deformation.

### Radiopacity

Radiopacity of the gel formulations was evaluated using microCT. One 2D image was rendered using a conventional high-pass threshold, while the second employed an ultra–high-pass threshold to assess attenuation at the upper limits of detection. Using a single global threshold, both the mouse femur and the K_2_HPO_4_ phantom could be readily segmented from all gel formulations, regardless of nanoparticle concentration or LT ratio. In both thresholding conditions, the gels reached the maximum value of the 16-bit grayscale range, indicating exceptionally high x-ray attenuation even at small feature sizes. Quantitative grayscale analysis further demonstrated the strong radiopaque performance of the gels. All gel formulations exhibited significantly higher radiopacity than mouse femur bone (p < 0.0001) and were comparable to or exceeded the attenuation of the 1,000 mg/mL K_2_HPO_4_ phantom.

## Discussion

This study demonstrates that nanoparticle-loaded alginate hydrogels incorporating lead abellaite and lead tungstate can meet the competing requirements of perfusability, mechanical robustness, long-term stability, and high radiopacity needed for post-mortem vascular microCT imaging. Rheological characterization showed that, in the absence of GDL, all formulations maintained low complex viscosity over extended time scales, with no evidence of shear-induced or spontaneous gelation. Although time, lead tungstate concentration, and LT–AB interactions influenced viscosity, the absolute values remained sufficiently low to support vascular perfusion without risk of blockage. This behavior is critical for consistent filling of small-diameter vessels and confirms that gelation can be reliably decoupled from the perfusion step.

Mechanical testing revealed that calcium exposure plays a dominant role in defining the functional strength of the gels. While traditional controls such as gelatin and Microfil tolerated larger overall strains before failure, the nanoparticle–alginate gels, particularly after calcium soaking, generated higher stresses within the low-strain regime most relevant to tissue handling. At strains of 0.1– 0.25, which approximate deformation experienced during organ manipulation and dissection, calcium-soaked gels consistently outperformed unsoaked gels and controls. Two-way ANOVA confirmed that calcium was the only factor significantly affecting mechanical properties across formulations, enhancing stiffness, elasticity, toughness, and stress at failure while reducing failure strain. Importantly, these mechanical gains directly support the intended application, as increased low-strain stiffness minimizes deformation of perfused vasculature during handling without rendering the material excessively brittle.

Long-term structural stability further distinguished calcium-treated gels from their unsoaked counterparts. Gels formed with GDL alone gradually lost structural integrity and reliquified over time, likely due to unstable lead–alginate complexation. In contrast, calcium-soaked gels remained macroscopically stable over extended storage periods, demonstrating that ionic crosslinking is essential not only for mechanical reinforcement but also for preserving gel architecture. This stability is particularly important for workflows that require delayed imaging, prolonged storage, or downstream histological processing.

Finally, radiopacity measurements confirmed that all gel formulations provide exceptionally strong x-ray attenuation. Independent of nanoparticle concentration or composition, the gels exhibited grayscale values significantly exceeding those of native bone and comparable to high-density calibration phantoms. The ability to segment bone, phantom, and gel using a single global threshold underscores the robustness of contrast generation and simplifies image processing. The fact that all formulations reached the upper limits of the 16-bit grayscale range further highlights their suitability for resolving fine vascular features without compromising dynamic range.

Taken together, these results indicate that calcium-stabilized, nanoparticle-loaded alginate gels offer a balanced combination of low-viscosity perfusion, mechanically appropriate stiffness for tissue handling, long-term stability, and superior radiographic contrast. By integrating controlled gelation with high intrinsic radiopacity, this system addresses key limitations of existing vascular contrast agents and provides a versatile platform for high-fidelity microCT angiography in orthopedic and broader biomedical research contexts.

## Conclusion

This study builds upon our previous development of an alginate-based vascular contrast agent by introducing a composite nanoparticle formulation incorporating both abellaite and lead tungstate. By leveraging the combined x-ray attenuation properties of lead and tungsten, this approach is designed to enhance overall radiopacity relative to abellaite alone. The inclusion of lead tungstate is expected to allow effective vascular contrast at reduced nanoparticle concentrations, which may improve injectability and uniform perfusion throughout the vascular network. Furthermore, the absence of a carbonate group in lead tungstate is anticipated to reduce gas generation during alginate gelation, thereby minimizing bubble-related imaging artifacts. Collectively, these improvements have the potential to enhance vessel continuity and image fidelity in post-mortem microCT angiography, particularly within small-diameter capillaries. This work provides a foundation for optimizing hydrogel-based, nanoparticle-enhanced contrast agents for high-resolution vascular imaging applications.

## Appendix

**Figure X:**
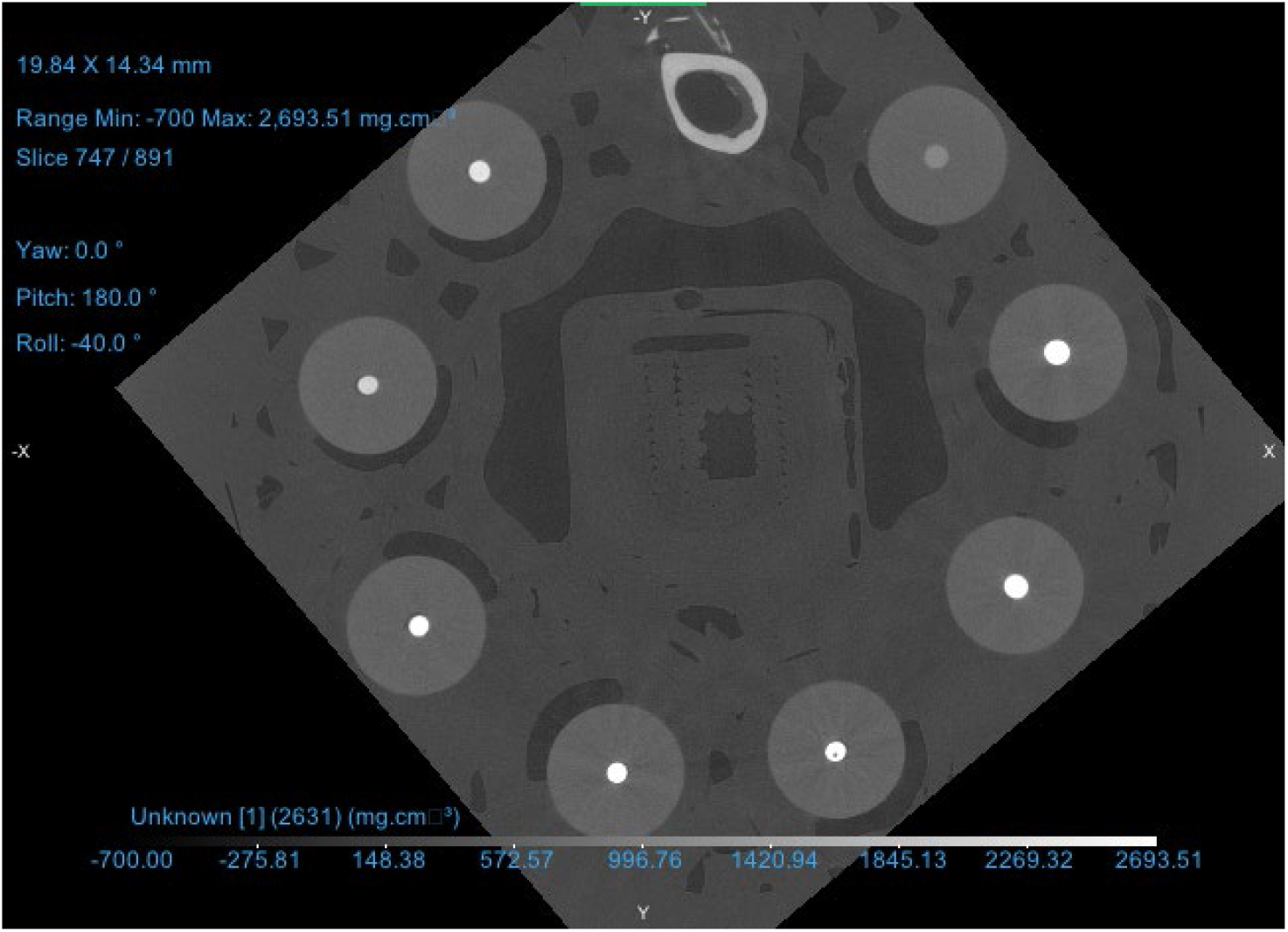
MicroCT X-Y view scan 5213

**Figure X:**
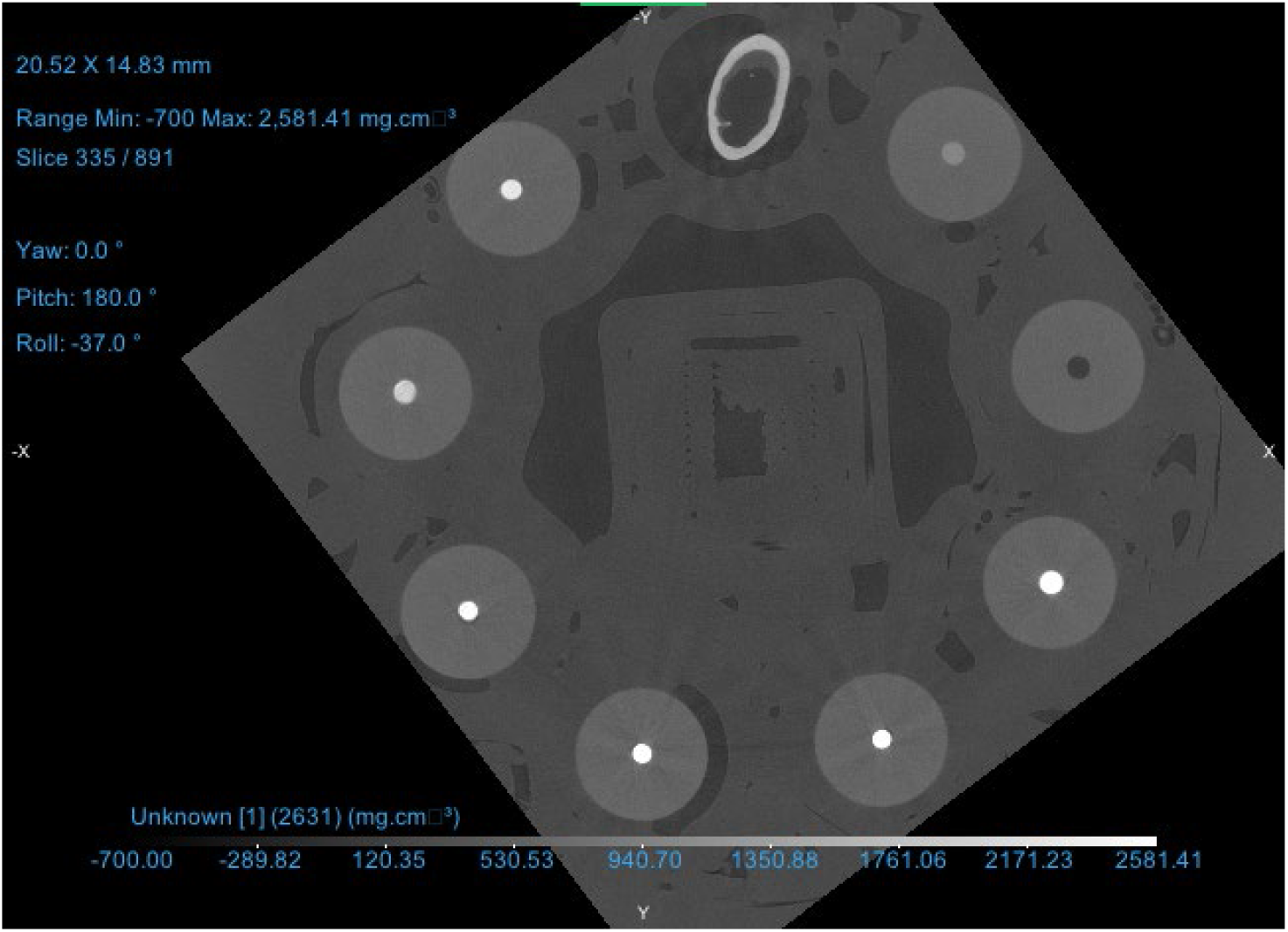
MicroCT X-Y view scan 5214

**Figure 9:**
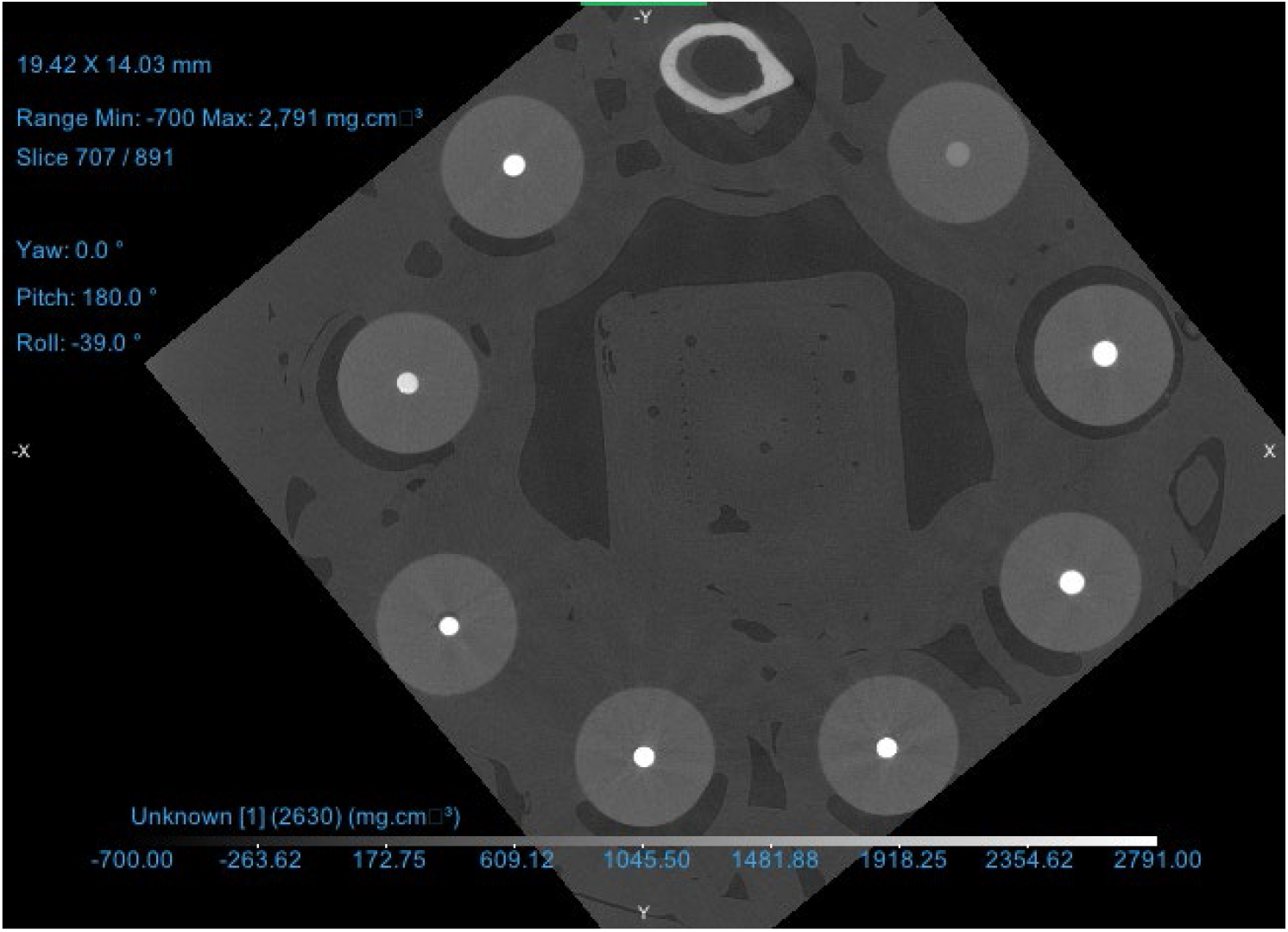
MicroCT X-Y view scan 5215

**Figure X:**
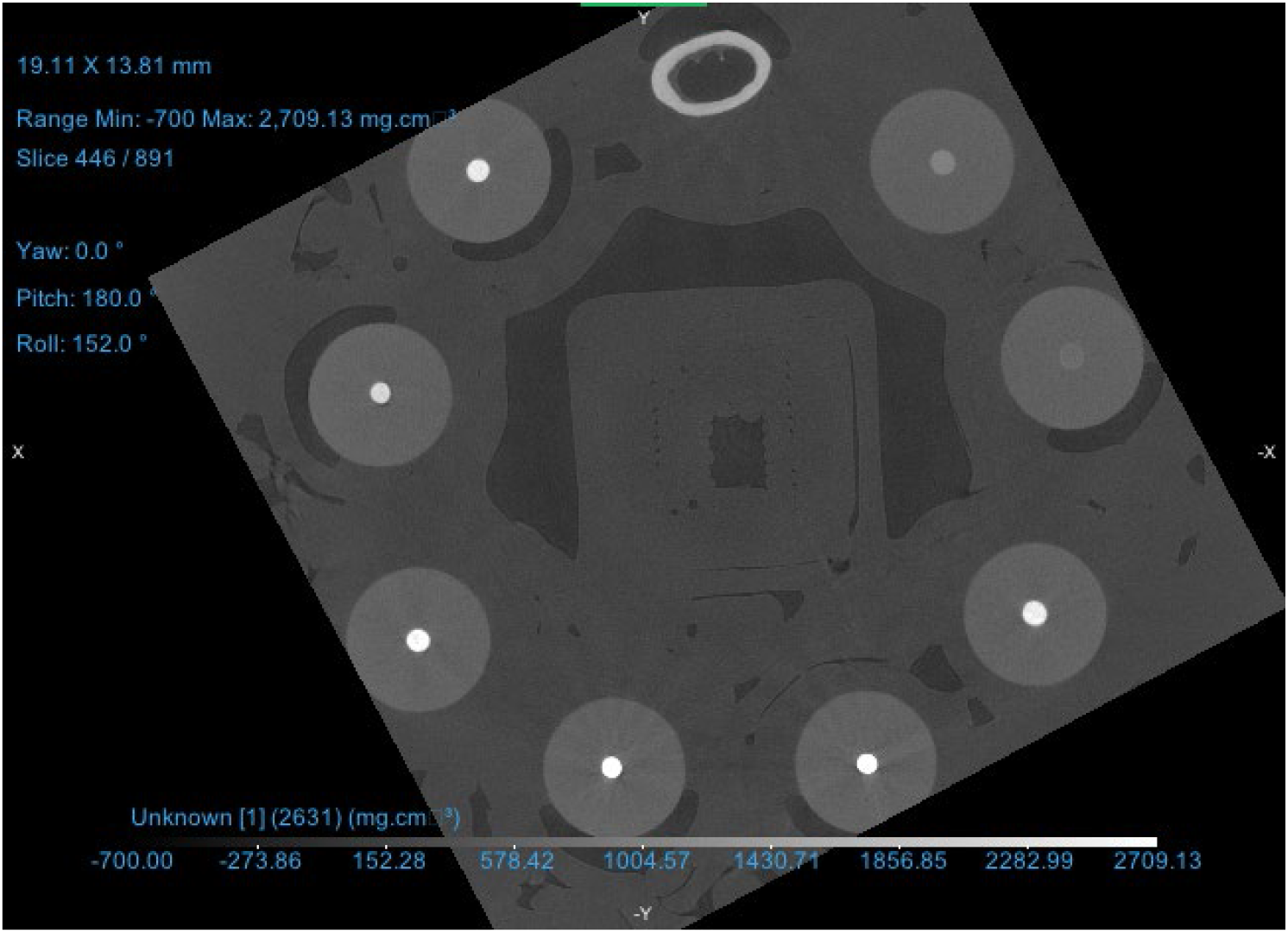
MicroCT X-Y view scan 5216

**Figure X:**
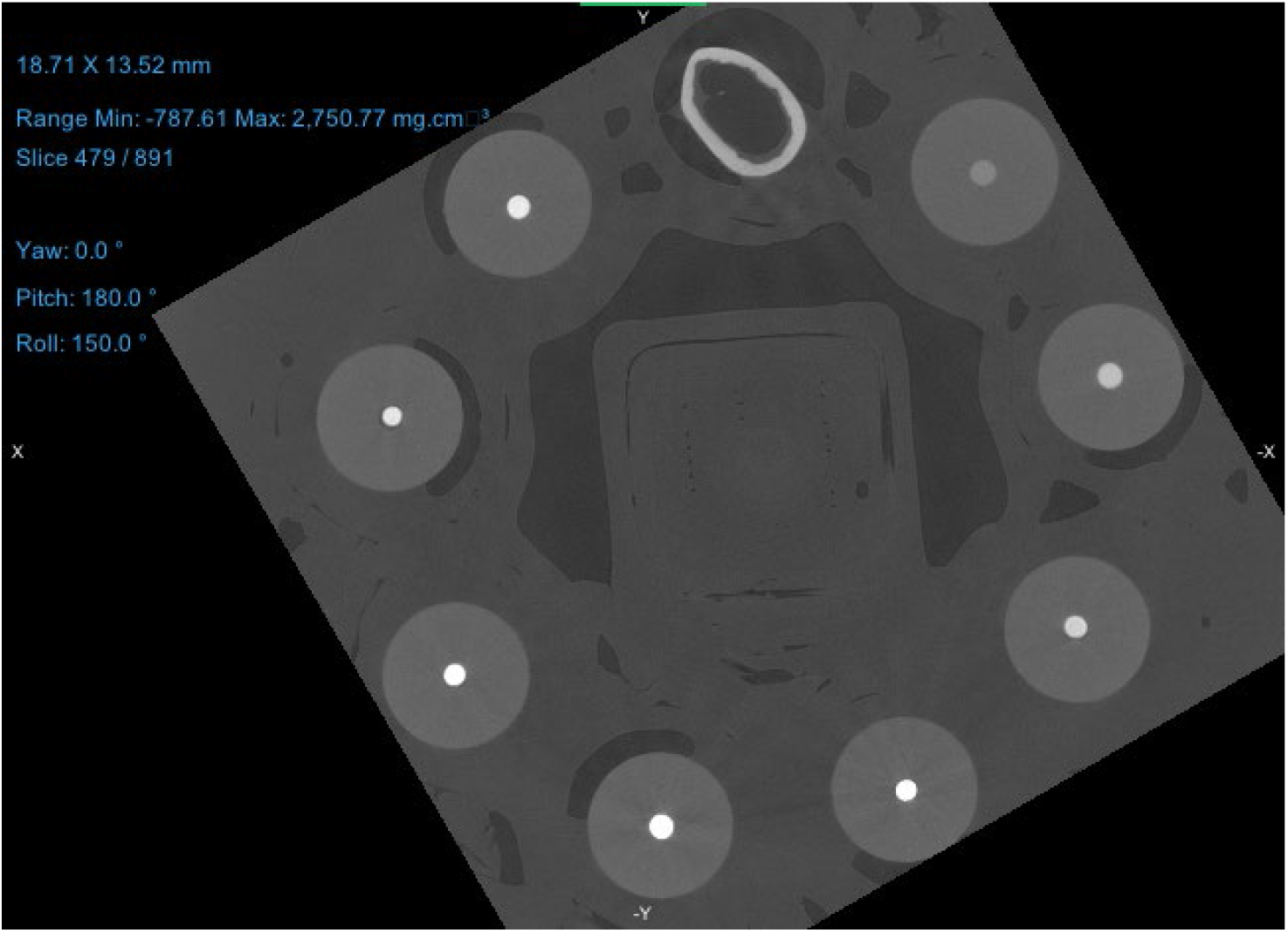
MicroCT X-Y view scan 5217

**Figure X:**
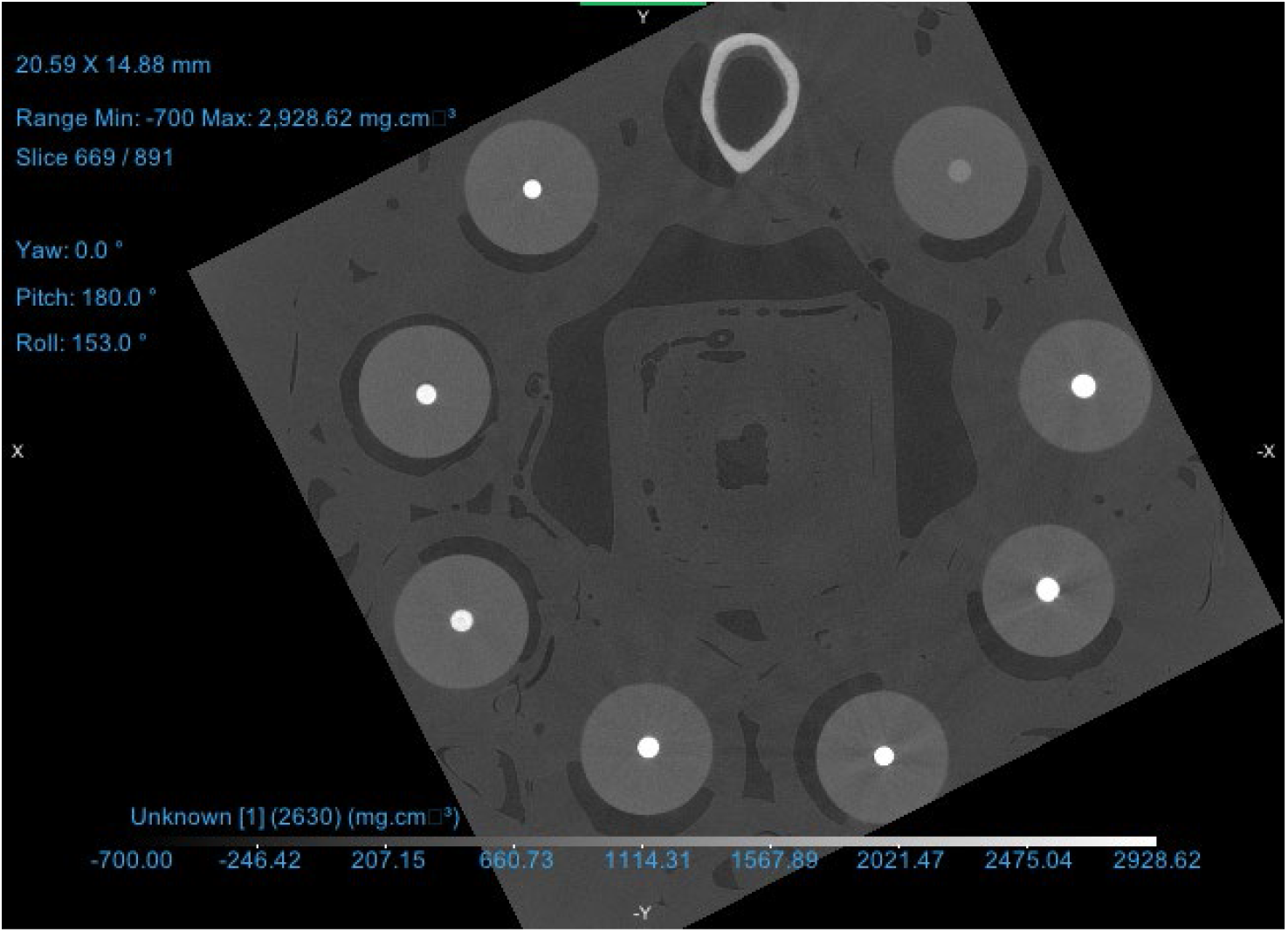
MicroCT X-Y view scan 5218

**Figure X:**
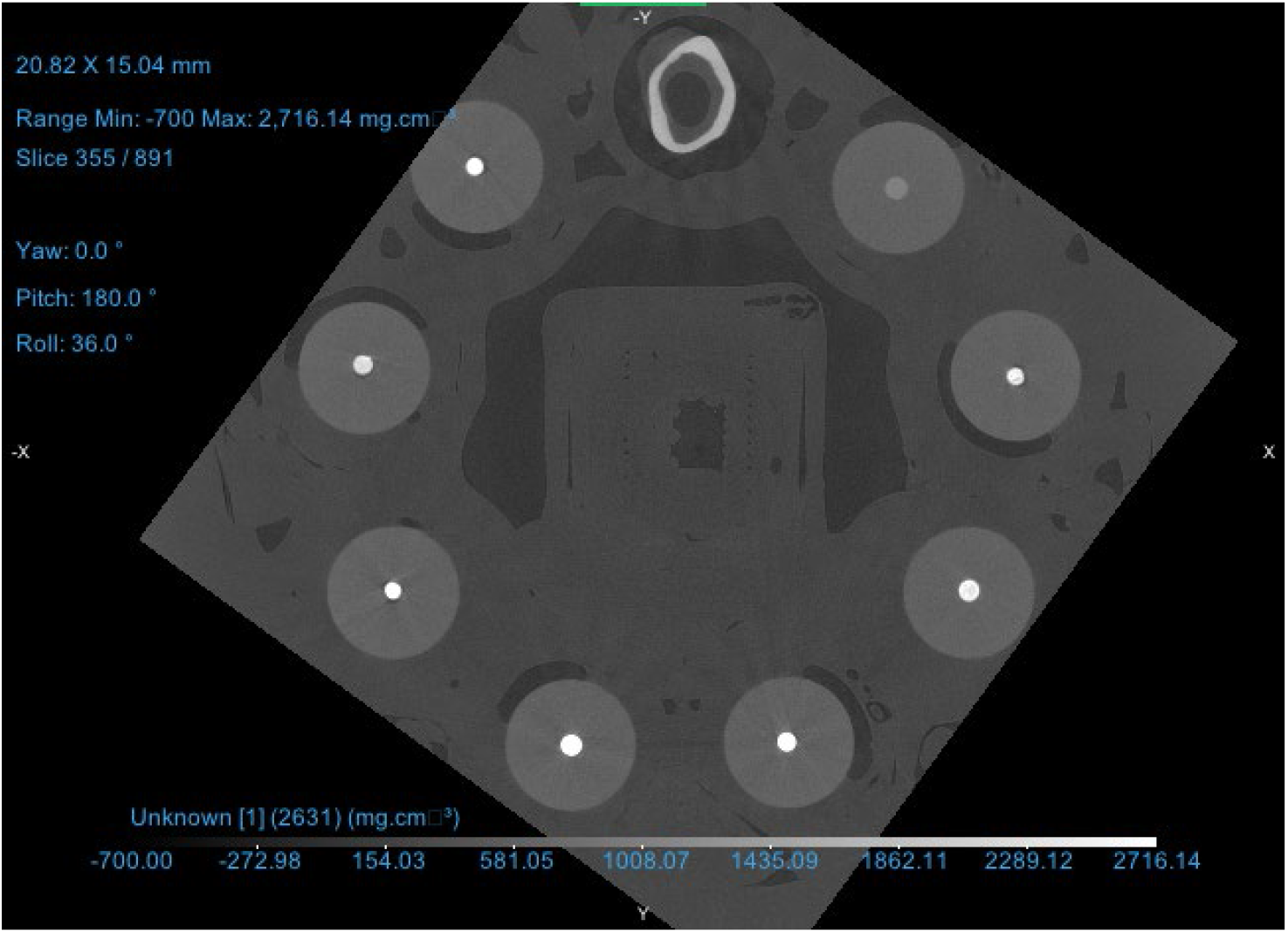
MicroCT X-Y view scan 5219

**Figure X:**
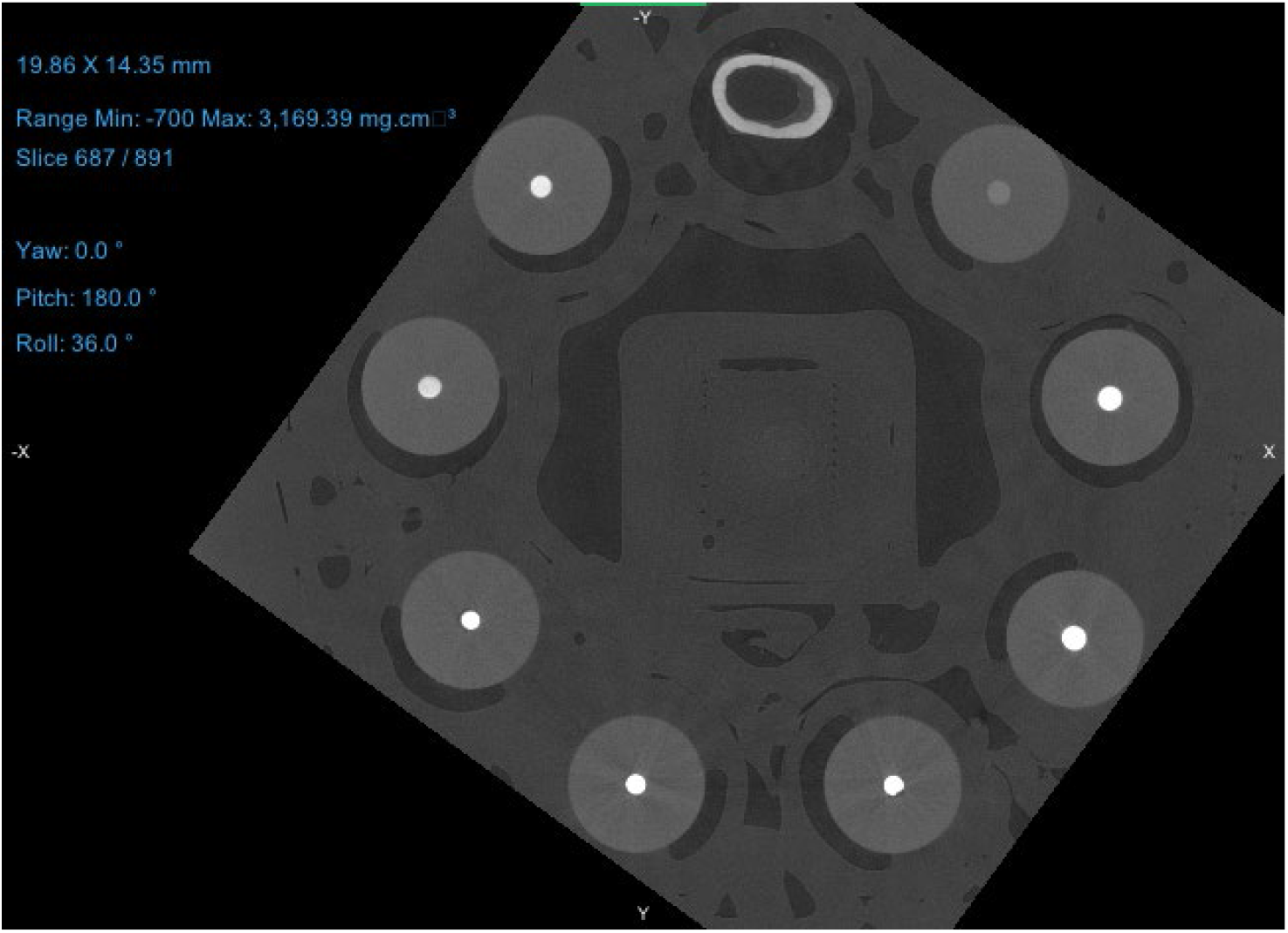
MicroCT X-Y view scan 5220

**Figure X:**
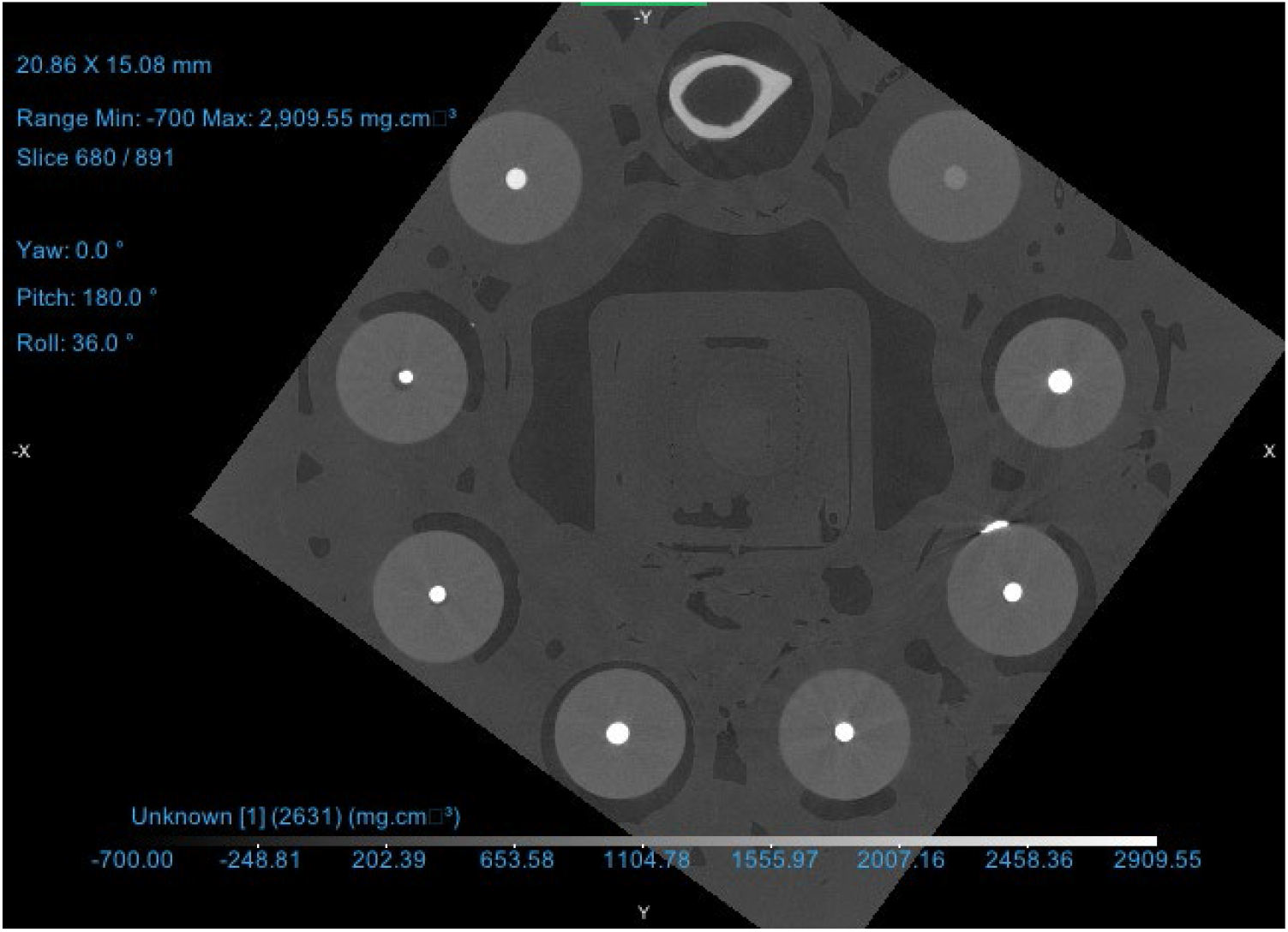
MicroCT X-Y view scan 5221

**Table X:** Bone Mineral Density Scan 5213.

| Name (NA) | Voxel count | Volume (mm<sup>3</sup>) | Min Intensity (mg.cm<sup>-3</sup>) | Max Intensity (mg.cm<sup>-3</sup>) | Mean Intensity (mg.cm<sup>-3</sup>) | Variance Intensity (mg.cm<sup>-3</sup>) | Std Dev Intensity (mg.cm<sup>-3</sup>) |
| --- | --- | --- | --- | --- | --- | --- | --- |
| Bone | 4844264 | 4.844264 | 1082.245 | 1726.643 | 1430.963 | 16531.09 | 128.5733 |
| Phantom | 519845 | 0.519845 | 777.6128 | 1030.186 | 928.8199 | 2880.001 | 53.66564 |
| 0.4 0:1 | 570997 | 0.570997 | 1734.759 | 2876.88 | 2562.796 | 112402.7 | 335.2651 |
| 0.4 1:1 | 728384 | 0.728384 | 1927.158 | 2876.88 | 2769.921 | 45951.14 | 214.3622 |
| 0.4 2:1 | 505592 | 0.505592 | 2155.78 | 2876.88 | 2852.431 | 11322.93 | 106.4092 |
| 0.4 5:1 | 533487 | 0.533487 | 2020.19 | 2876.88 | 2845.913 | 17109.47 | 130.8032 |
| 0.2 0:1 | 484031 | 0.484031 | 1957.245 | 2876.88 | 2702.066 | 47083.09 | 216.9864 |
| 0.2 1:1 | 447158 | 0.447158 | 1420.625 | 2141.429 | 1870.082 | 27860.47 | 166.9145 |
| 0.2 2:1 | 508031 | 0.508031 | 1532.264 | 2482.977 | 2195.896 | 42336.36 | 205.758 |

**Table X:** Bone Mineral Density Scan 5214.

| Name (NA) | Voxel count | Volume (mm<sup>3</sup>) | Min Intensity (mg.cm<sup>-3</sup>) | Max Intensity (mg.cm<sup>-3</sup>) | Mean Intensity (mg.cm<sup>-3</sup>) | Variance Intensity (mg.cm<sup>-3</sup>) | Std Dev Intensity (mg.cm<sup>-3</sup>) |
| --- | --- | --- | --- | --- | --- | --- | --- |
| Bone | 4650860 | 4.65086 | 1145.343 | 1788.356 | 1546.837 | 15907.35 | 126.1243 |
| Phantom | 482674 | 0.482674 | 751.8906 | 1072.516 | 956.5354 | 4375.601 | 66.14833 |
| 0.4 1:1 | 685959 | 0.685959 | 2130.944 | 3026.002 | 2974.448 | 27720.38 | 166.4944 |
| 0.4 2:1 | 514546 | 0.514546 | 2038.537 | 3026.002 | 2985.171 | 25959.03 | 161.1181 |
| 0.4 5:1 | 482068 | 0.482068 | 2076.66 | 3026.002 | 2989.371 | 22601.11 | 150.3367 |
| 0.2 0:1 | 459823 | 0.459823 | 2043.924 | 3026.002 | 2913.045 | 46287.71 | 215.1458 |
| 0.2 1:1 | 629956 | 0.629956 | 1386.305 | 2339.376 | 2027.507 | 46135.36 | 214.7914 |
| 0.2 2:1 | 390682 | 0.390682 | 1662.696 | 2579.405 | 2230.938 | 40591.05 | 201.4722 |

**Table X:** Bone Mineral Density Scan 5215.

| Name (NA) | Voxel count | Volume (mm<sup>3</sup>) | Min Intensity (mg.cm<sup>-3</sup>) | Max Intensity (mg.cm<sup>-3</sup>) | Mean Intensity (mg.cm<sup>-3</sup>) | Variance Intensity (mg.cm<sup>-3</sup>) | Std Dev Intensity (mg.cm<sup>-3</sup>) |
| --- | --- | --- | --- | --- | --- | --- | --- |
| Bone | 4341926 | 4.341926 | 1326.929 | 1893.894 | 1613.735 | 15359.15 | 123.932 |
| Phantom | 584101 | 0.584101 | 798.1672 | 1155.46 | 1013.675 | 5113.643 | 71.50974 |
| 0.4 1:1 | 669640 | 0.66964 | 1757.646 | 3246.55 | 2859.665 | 159340.6 | 399.1749 |
| 0.4 2:1 | 487185 | 0.487185 | 2229.325 | 3246.55 | 3205.429 | 26646.69 | 163.2381 |
| 0.4 5:1 | 540275 | 0.540275 | 2356.078 | 3246.55 | 3215.261 | 17952.47 | 133.9868 |
| 0.2 0:1 | 459515 | 0.459515 | 1878.547 | 3246.55 | 3039.532 | 115668.7 | 340.101 |
| 0.2 1:1 | 463804 | 0.463804 | 2085.238 | 3246.55 | 2613.846 | 108028 | 328.6761 |
| 0.2 2:1 | 585692 | 0.585692 | 1985.867 | 3246.55 | 2806.2 | 105333.9 | 324.5519 |

**Table X:**
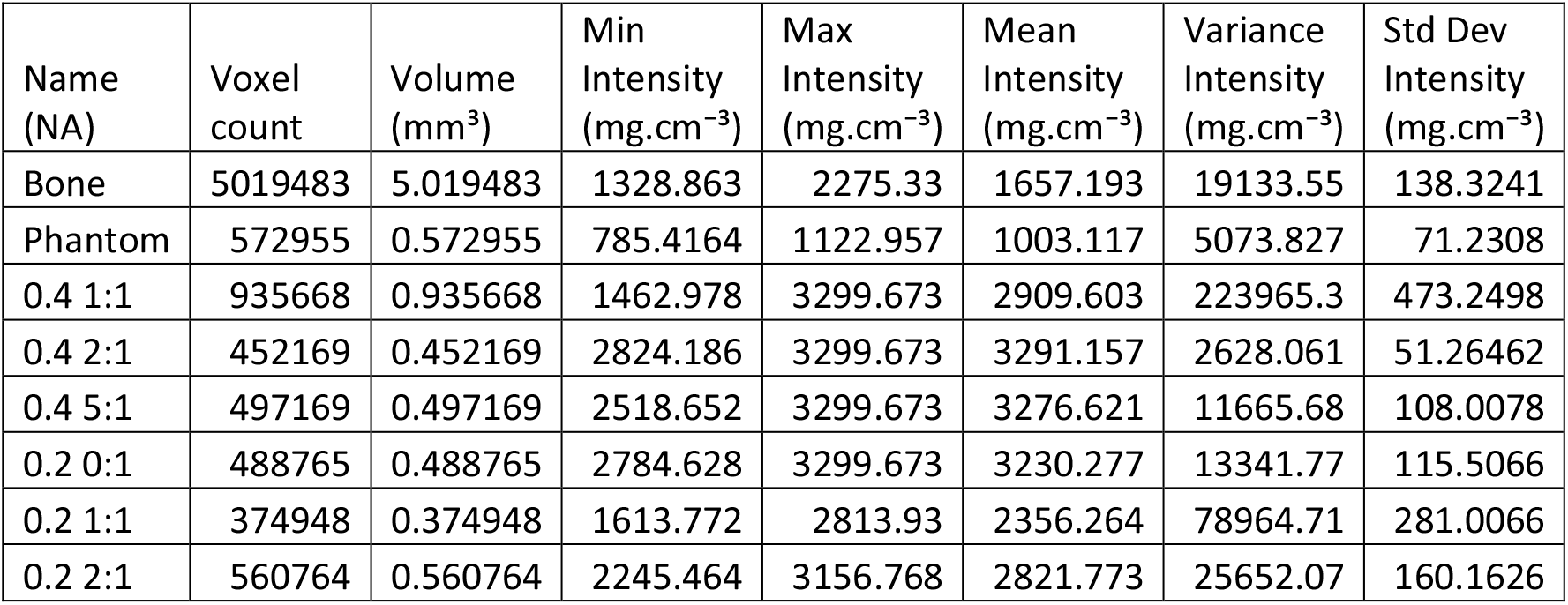
Bone Mineral Density Scan 5216.

**Table X:** Bone Mineral Density Scan 5217.

| Name (NA) | Voxel count | Volume (mm<sup>3</sup>) | Min Intensity (mg.cm<sup>-3</sup>) | Max Intensity (mg.cm<sup>-3</sup>) | Mean Intensity (mg.cm<sup>-3</sup>) | Variance Intensity (mg.cm<sup>-3</sup>) | Std Dev Intensity (mg.cm<sup>-3</sup>) |
| --- | --- | --- | --- | --- | --- | --- | --- |
| Bone | 7025755 | 7.025755 | 464.7208 | 1915.33 | 1450.805 | 141672 | 376.3934 |
| Phantom | 571911 | 0.571911 | 765.0464 | 1107.466 | 981.3727 | 5323.909 | 72.96512 |
| 0.4 0:1 | 302279 | 0.302279 | 3034.133 | 3449.524 | 3406.463 | 10025.83 | 100.1291 |
| 0.4 1:1 | 460422 | 0.460422 | 3222.048 | 3449.524 | 3442.32 | 944.0491 | 30.72538 |
| 0.4 2:1 | 521531 | 0.521531 | 2222.41 | 3449.524 | 3394.173 | 43095.24 | 207.5939 |
| 0.4 5:1 | 740859 | 0.740859 | 2019.539 | 3449.524 | 3365.468 | 75228.29 | 274.2778 |
| 0.2 0:1 | 407724 | 0.407724 | 2494.874 | 3449.524 | 3391.402 | 33111.35 | 181.9652 |
| 0.2 1:1 | 496243 | 0.496243 | 2190.086 | 3449.524 | 3055.462 | 127010.7 | 356.3856 |
| 0.2 2:1 | 477341 | 0.477341 | 1936.316 | 3449.524 | 2871.981 | 89076.15 | 298.4563 |

**Table X:** Bone Mineral Density Scan 5218.

| Name (NA) | Voxel count | Volume (mm<sup>3</sup>) | Min Intensity (mg.cm<sup>-3</sup>) | Max Intensity (mg.cm<sup>-3</sup>) | Mean Intensity (mg.cm<sup>-3</sup>) | Variance Intensity (mg.cm<sup>-3</sup>) | Std Dev Intensity (mg.cm<sup>-3</sup>) |
| --- | --- | --- | --- | --- | --- | --- | --- |
| Bone | 4685015 | 4.685015 | 1519.579 | 2167.041 | 1731.188 | 14558.89 | 120.6602 |
| Phantom | 503154 | 0.503154 | 816.9881 | 1142.671 | 1025.67 | 5125.076 | 71.58964 |
| 0.4 0:1 | 289194 | 0.289194 | 1485.981 | 3460.783 | 3301.195 | 234842.3 | 484.6053 |
| 0.4 1:1 | 678806 | 0.678806 | 2436.886 | 3460.783 | 3375.413 | 49993.26 | 223.5917 |
| 0.4 2:1 | 503729 | 0.503729 | 2242.517 | 3460.783 | 3402.052 | 45700.38 | 213.7765 |
| 0.4 5:1 | 579217 | 0.579217 | 2436.177 | 3460.783 | 3422.872 | 24888 | 157.7593 |
| 0.2 0:1 | 544521 | 0.544521 | 2236.957 | 3460.783 | 3128.922 | 81832.26 | 286.0634 |
| 0.2 1:1 | 444578 | 0.444578 | 2719.153 | 3460.783 | 3213.856 | 46958.55 | 216.6992 |
| 0.2 2:1 | 450600 | 0.4506 | 2553.531 | 3460.783 | 3388.648 | 34714.96 | 186.3195 |

**Table X:** Bone Mineral Density Scan 5219.

| Name (NA) | Voxel count | Volume (mm<sup>3</sup>) | Min Intensity (mg.cm<sup>-3</sup>) | Max Intensity (mg.cm<sup>-3</sup>) | Mean Intensity (mg.cm<sup>-3</sup>) | Variance Intensity (mg.cm<sup>-3</sup>) | Std Dev Intensity (mg.cm<sup>-3</sup>) |
| --- | --- | --- | --- | --- | --- | --- | --- |
| Bone | 5144399 | 5.144399 | 1339.471 | 2152.197 | 1736.193 | 20569.42 | 143.4204 |
| Phantom | 476390 | 0.47639 | 647.3604 | 1238.084 | 998.4586 | 5045.765 | 71.03355 |
| 0.4 0:1 | 444388 | 0.444388 | 1612.166 | 3391.097 | 2979.612 | 283366.6 | 532.3219 |
| 0.4 1:1 | 763675 | 0.763675 | 2139.727 | 3391.097 | 3118.004 | 115886.9 | 340.4216 |
| 0.4 2:1 | 493155 | 0.493155 | 2470.108 | 3391.097 | 3359.682 | 18791.25 | 137.0812 |
| 0.4 5:1 | 605506 | 0.605506 | 2519.287 | 3391.097 | 3362.256 | 16078.95 | 126.8028 |
| 0.2 0:1 | 366446 | 0.366446 | 2566.134 | 3391.097 | 3351.092 | 20352.72 | 142.663 |
| 0.2 1:1 | 257294 | 0.257294 | 1879.152 | 3068.757 | 2689.437 | 95232.61 | 308.5978 |
| 0.2 2:1 | 442208 | 0.442208 | 2261.974 | 3391.097 | 3306.064 | 55705.52 | 236.0202 |

**Table X:**
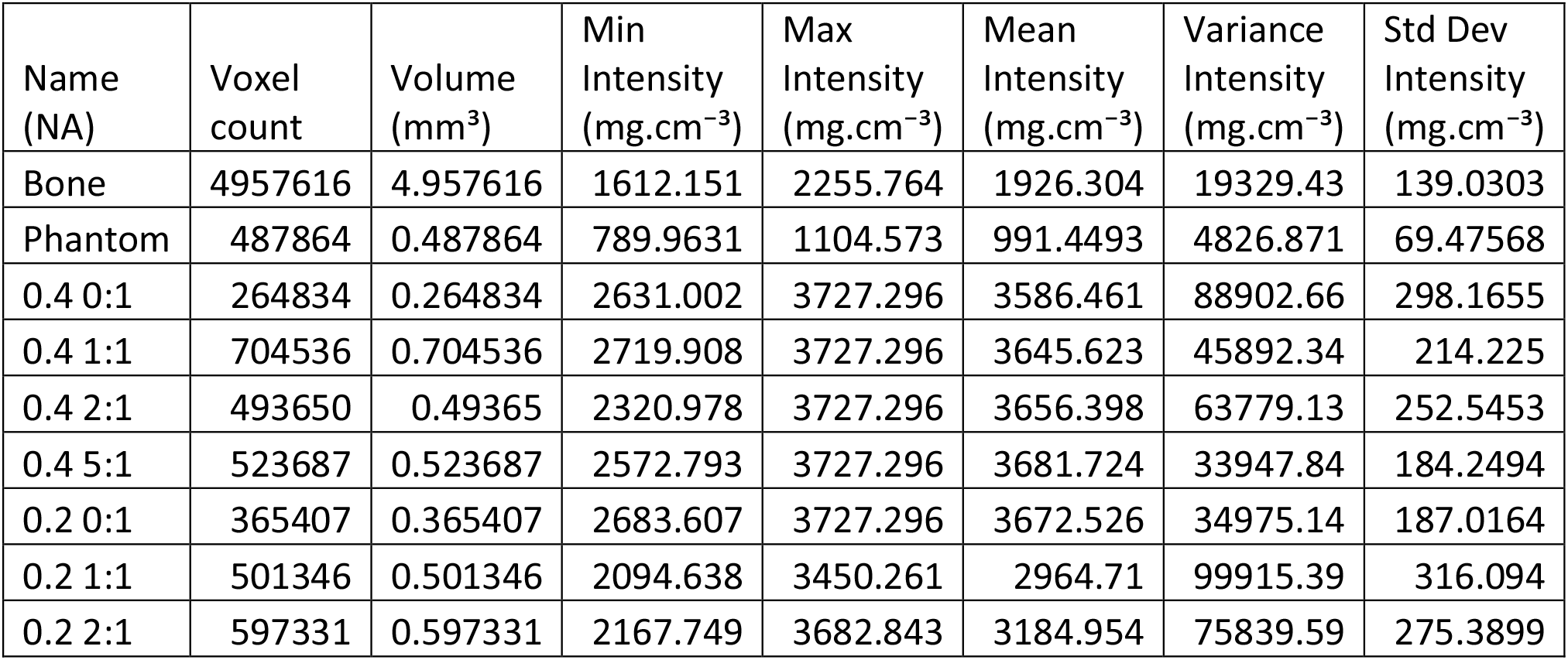
Bone Mineral Density Scan 5220.

**Table X:** Bone Mineral Density Scan 5221.

| Name (NA) | Voxel count | Volume (mm<sup>3</sup>) | Min Intensity (mg.cm<sup>-3</sup>) | Max Intensity (mg.cm<sup>-3</sup>) | Mean Intensity (mg.cm<sup>-3</sup>) | Variance Intensity (mg.cm<sup>-3</sup>) | Std Dev Intensity (mg.cm<sup>-3</sup>) |
| --- | --- | --- | --- | --- | --- | --- | --- |
| Bone | 5742752 | 5.742752 | 764.4693 | 2093.52 | 1592.332 | 66698.9 | 258.2613 |
| Phantom | 664230 | 0.66423 | 751.5084 | 1134.525 | 949.0234 | 5116.281 | 71.52819 |
| 0.4 0:1 | 213547 | 0.213547 | 2916.76 | 3278.883 | 3260.501 | 4170.889 | 64.58242 |
| 0.4 1:1 | 507936 | 0.507936 | 2177.542 | 3278.883 | 3198.572 | 50855.75 | 225.5122 |
| 0.4 2:1 | 461961 | 0.461961 | 2818.994 | 3278.883 | 3270.838 | 2426.031 | 49.25476 |
| 0.4 5:1 | 603205 | 0.603205 | 2609.721 | 3278.883 | 3258.818 | 8559.44 | 92.51724 |
| 0.2 0:1 | 343149 | 0.343149 | 2529.609 | 3278.883 | 3245.99 | 15384.36 | 124.0337 |
| 0.2 1:1 | 300453 | 0.300453 | 2058.547 | 3278.883 | 3190.305 | 65870.18 | 256.6519 |
| 0.2 2:1 | 614405 | 0.614405 | 1910.391 | 3252.514 | 2764.256 | 54675.36 | 233.8276 |

## References

1. Sheikh, A. Md., Yano, S., Tabassum, S. & Nagai, A. The Role of the Vascular System in Degenerative Diseases: Mechanisms and Implications. Int J Mol Sci 25, 2169 (2024).

2. Ghadimi, M. & Thomas, A. Magnetic Resonance Imaging Contraindications. in StatPearls (StatPearls Publishing, Treasure Island (FL), 2025).

3. Mantella, L. E., Liblik, K. & Johri, A. M. Vascular imaging of atherosclerosis: Strengths and weaknesses. Atherosclerosis 319, 42–50 (2021).

4. Tiwari, A., Elgrably, B., Saar, G. & Vandoorne, K. Multi-Scale Imaging of Vascular Pathologies in Cardiovascular Disease. Front Med (Lausanne) 8, 754369 (2022).

5. Singh, V. & Sandean, D. P. CT Patient Safety And Care. in StatPearls (StatPearls Publishing, Treasure Island (FL), 2025).

6. Florkow, M. C. et al. Magnetic Resonance Imaging Versus Computed Tomography for Three-Dimensional Bone Imaging of Musculoskeletal Pathologies: A Review. J Magn Reson Imaging 56, 11–34 (2022).

7. van Beek, E. J. R. et al. Value of MRI in Medicine: More Than Just Another Test? J Magn Reson Imaging 49, e14–e25 (2019).

8. Tu, L. H. et al. Cost-Effectiveness of CT, CTA, MRI, and Specialized MRI for Evaluation of Patients Presenting to the Emergency Department With Dizziness. American Journal of Roentgenology 222, e2330060 (2024).

9. X-Ray Mass Attenuation Coefficients. NIST https://www.nist.gov/pml/x-ray-mass-attenuation-coefficients (2009).

10. Grabherr, S., Grimm, J., Baumann, P. & Mangin, P. Application of contrast media in post-mortem imaging (CT and MRI). Radiol med 120, 824–834 (2015).

11. Kader, M. S. et al. Synthesis and Characterization of BaSO4–CaCO3–Alginate Nanocomposite Materials as Contrast Agents for Fine Vascular Imaging. ACS Mater. Au 2, 260–268 (2022).

12. Koç, M. M., Aslan, N., Kao, A. P. & Barber, A. H. Evaluation of X-ray tomography contrast agents: A review of production, protocols, and biological applications. Microscopy Research and Technique 82, 812–848 (2019).

13. Margolis, R. et al. Comparison of micro-CT image enhancement after use of different vascular casting agents. Quant Imaging Med Surg 14, 2568–2579 (2024).

14. Le, A. X. et al. A Lead Abellaite-Alginate Nanocomposite as a Contrast Agent for Post-Mortem CT Imaging.

15. Gamage, M. E. et al. Synthesis of Lead(II) Carbonate-Containing Nanoparticles Using Ultrasonication or Microwave Irradiation. ACS Omega 9, 48802–48809 (2024).

16. Hubbell, J. H. & Seltzer, S. M. Tables of X-Ray Mass Attenuation Coefficients and Mass Energy-Absorption Coefficients 1 keV to 20 MeV for Elements Z = 1 to 92 and 48 Additional Substances of Dosimetric Interest. NIST https://www.nist.gov/publications/tables-x-ray-mass-attenuation-coefficients-and-mass-energy-absorption-coefficients-1-0 (1995).

